# A functional hierarchy for choice in medial prefrontal cortex

**DOI:** 10.1101/2020.07.28.226043

**Authors:** David J-N. Maisson, Tyler V. Cash-Padgett, Benjamin Y. Hayden, Sarah R. Heilbronner, Jan Zimmermann

**Affiliations:** Department of Neuroscience, Center for Magnetic Resonance Research, Center for Neuroengineering Department of Biomedical Engineering University of Minnesota, Minneapolis MN 55455

**Keywords:** anterior cingulate cortex, medial wall, pregenual, subgenual, ventromedial prefrontal cortex

## Abstract

Hierarchical approaches to functional neuroanatomy propose that choice-relevant brain regions have overlapping functions and can be organized into a series that progressively transforms information about options into choices. Here, we examined responses of neurons in four regions of the medial prefrontal cortex as macaques performed two-option risky choices. All four regions encoded economic variables in similar proportions and showed putative signatures of key choice-related computations. We found evidence for a hierarchical organization proceeding from areas 14→25→32→24. Specifically, we found that decodability of eight distinct task variables increased along that path, consistent with the idea that hierarchically later regions make these variables more separable. We also found longer intrinsic timescales in the same series, further supporting the idea of a hierarchy. Together these results highlight the importance of the medial wall in choice, endorse a specific hierarchical organization, and argue against a modular functional neuroanatomy of choice.

## INTRODUCTION

Economic choice is mediated by a large number of brain regions, including several in the prefrontal cortex (Rangel et al., 2008; Haber and Knutson, 2010; Rushworth et al., 2011; Haber and Beherens, 2014). The principles delineating the parcellation of function in choice remain unclear. On one hand, modular theories hold that specific brain regions can be associated with particular nameable functions, such as evaluation, comparison, and action selection (Rangel et al., 2008; Kable and Glimcher, 2009; Padoa-Schioppa, 2011; Passingham and Wise, 2015; Hauser et al., 2015; Hunt et al., 2018). On the other hand, distributed approaches to understanding choice hold that particular elements of choice do not correspond neatly to anatomical regions (Cisek and Kalaska, 2010; Cisek, 2012; Hunt and Hayden, 2017; Yoo and Hayden, 2018). Distributed approaches are inspired by classic connectionist theories as well as by modern deep learning approaches (Plaut, 1995; Bengio et al., 2003; Pennington et al., 2014; Hunt and Hayden, 2017; Balasubramani et al., 2018). They are also inspired by cognitive and philosophical theories of distributed cognition (Uttal et al., 2001; Prinz, 2006; Anderson, 2007), and by analogy to the form vision system, where hierarchical models have come to replace historically dominant modular models (Maunsell and van Essen, 1983; DiCarlo et al., 2012; Yoo and Hayden, 2018).

Within the domain of distributed models, early mass-action and equipotentiality models have lately been supplanted by *hierarchical distributed* models (Fuster, 1990; Fellemen and Van Essen, 1991; Rushworth et al., 2011). We and others have extended the approach by proposing one described by a gradual transformation of information, but that networked areas show graded functional distinctions (Hunt and Hayden, 2017; Hunt et al., 2012; Hunt et al., 2015). For example, specific circuits within the prefrontal cortex may be organized into a hierarchical gradient, so that each anatomical region implements part of a transformation of task-relevant information from a domain of options to a domain of actions. We have proposed that each region *untangles* information about the best action, which is latent in the early representations and which, through serial processing, is transformed into appropriate choice actions (Yoo and Hayden, 2018). This view is inspired by, and is an extension of, modern untangling-based models of form vision (DiCarlo and Cox, 2007; DiCarlo et al., 2012).

A critical prediction of hierarchical models is that it should be possible to arrange medial prefrontal regions into a particular hierarchical organization on the basis of their functional properties. Discussions of prefrontal hierarchy have typically focused on the lateral surface, or, when examining the medial wall, on the rostrocaudal axis (Koechlin et al., 2002; Rushworth et al., 2007; Badre, 2008; Nee et al., 2013; Fuster, 2001; Cai and Padoa-Schioppa, 2014; Siegel et al., 2015). We were interested, instead, in the ventrodorsal dimension of the medial prefrontal cortex. Neuroeconomists have long proposed that the orbitofrontal cortex serves as the entry point of choice-relevant sensory information into the prefrontal cortex and that the motor cortex serves as the output (e.g. Hare et al., 2011; Padoa-Schioppa, 2011; Rudebeck and Murray, 2014; Murray et al., 2014). The medial wall inferior to the premotor cortex, which includes areas 14, 25, 32, and 24, looks to be a likely pathway linking OFC to pre/motor areas. These regions also have prominent limbic, visceral, and reward-related functions, suggesting they may contribute to valuation and perhaps to choice (Vogt et al., 1979; Carmichael and Price, 1995, 1996; Freedman et al., 2000).

However, there are several possible functional hierarchies that are consistent with known anatomy. First, it could follow topology in a rough ventrodorsal direction (14→25→32→24). Second, it could match the contour of the genu of the corpus callosum (25→14→32→24). Third, cytoarchitecture suggests that the less differentiated cingulate areas (25, 32, and 24) may precede the more differentiated area 14 (Barbas and Pandya, 1989). Other cytoarchitectural studies suggest that pre- and subgenual regions (14, 25, and 32) may have shared function but differ qualitatively from the postgenual 24 (Bush et al., 2000). These specific pathways have not, to our knowledge, been functionally evaluated. Despite this, identifying the functional hierarchy, if one exists, is critical for establishing a neuroscience of economic choice (Rangel et al., 2008).

We examined a composite dataset consisting of single unit responses collected in these four brain regions, including previously published data for 14, 25, and 24, and newly collected data for area 32 (Strait et al., 2014; Azab and Hayden, 2017, 2018). Instead of looking for specific functions that would distinguish these regions from each other, we took a function-first approach: we selected key putative signatures of participation in specific elements of choice, and then characterized each region in these functions. Overall, we found two major results. First, the regions all show signatures of all tested functions and do not differ qualitatively in whether they carry certain information or have signatures of choice processes. Second, a decoding analysis shows both stronger decodability on eight dimensions consistent with a single hierarchy, specifically one that progresses from 14→25→32→24. This second result is complemented by a demonstration that intrinsic timescale shows the same pattern. Together these results support a specific ventrodorsal medial prefrontal hierarchy and, simultaneously, argue against a modular view in which conceptually distinct functions are reified in neuroanatomy.

## RESULTS

### Behavioral results

Rhesus macaque (*Macaca mulatta*) subjects performed one of two structurally similar two-alternative forced choice gambling tasks (Strait et al., 2014; Azab and Hayden, 2017; see **Methods**). Briefly, subjects chose between two risky options presented asynchronously (**Figure 1A-B**). After the second offer was presented, a go cue indicates that the subject was free to shift gaze toward the target option to indicate a choice. Thus, the task was naturally divided into three epochs, corresponding to the periods immediately following offer 1, offer 2, and choice. The offer 2 epoch is the first during which the subject can compare subjective values and select an action. (Note, however, that subjects can and likely do form tentative partial choices based solely on the value of the first offer, Azab and Hayden, 2017).

**Figure 1.**
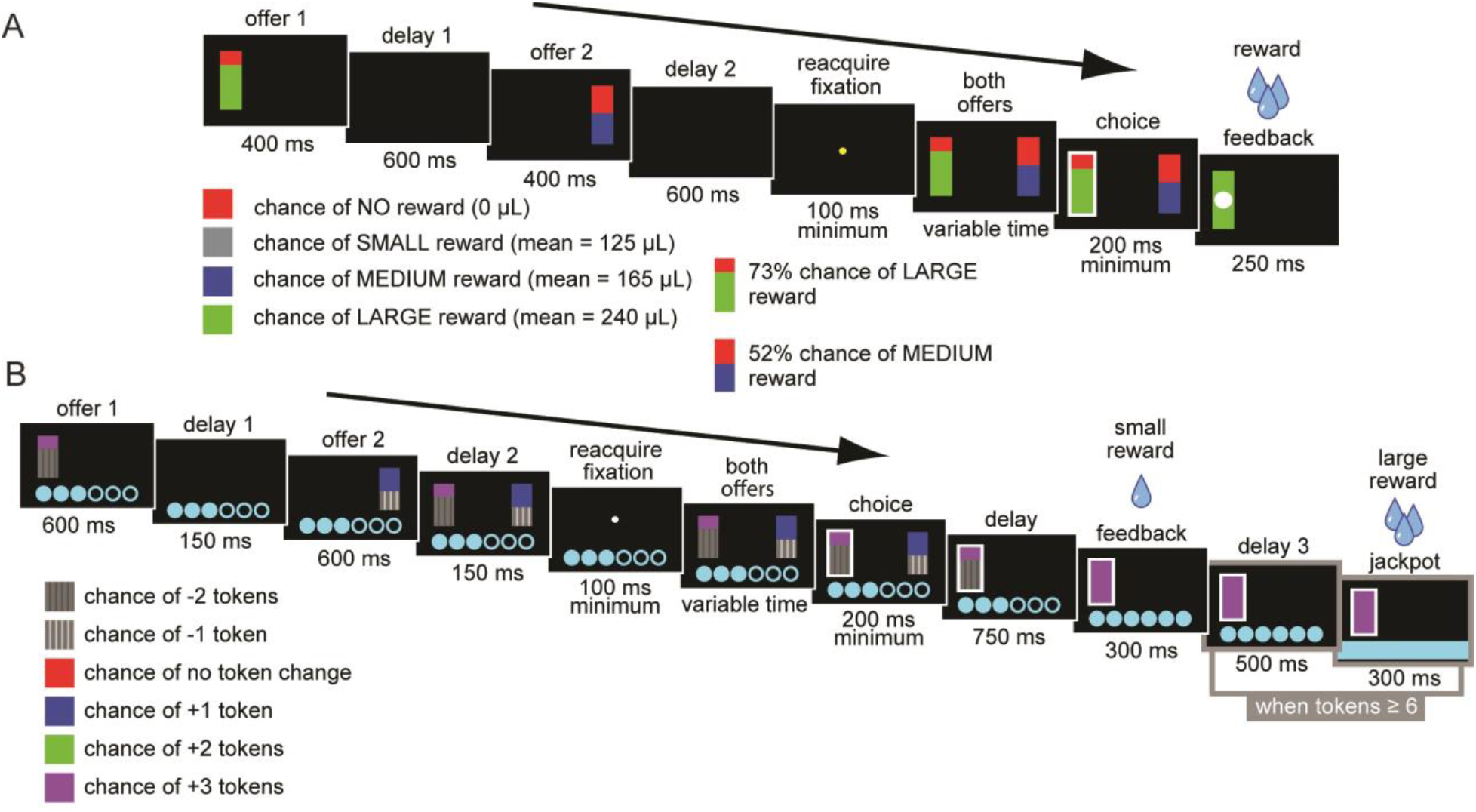
Gambling Tasks. A. Risky choice task: The first offer is presented, followed by a delay period, after which the second offer is presented. After another delay period, fixation is reacquired for a minimum of 100ms. Both offers are then presented, a choice is made, and the choice is probabilistically rewarded. Offers consist of a reward magnitude (color of the non-red portion of the bar) and a probability (size of the colored portion). B. Token risky choice task: Equivalent format but reward is tokenized. Once tokens reach 6, a reward is delivered.

Behavior in these tasks has been explored at length and is not reanalyzed here (for the most detailed analyses, see Farashahi et al., 2018; Farashahi et al., 2019). Briefly, behavior reflected understanding of all important task variables with very weak order or side biases. We defined the *expected value* of an offer as the product of the offer magnitude (in uL juice) and probability of reward. Thus, for a basic characterization of behavior, we computed the frequency with which a given offer was chosen when it had a higher expected value. We determined the proportion of trials on which the subject chose the first offer and we compared it to the difference in expected values of the two offers for each trial. Subjects’ behavior described a sigmoidal function (**Figure 2A**). Subjects most frequently chose the offer with the higher expected value (vmPFC sessions: 84.55% of trials; sgACC sessions: 78.62%; pgACC sessions: 74.97%; dACC sessions: 75.57%; *p* < 0.0001 in all cases; 1-sample t-test), consistent with the idea that they had a basic understanding of the task.

**Figure 2.**
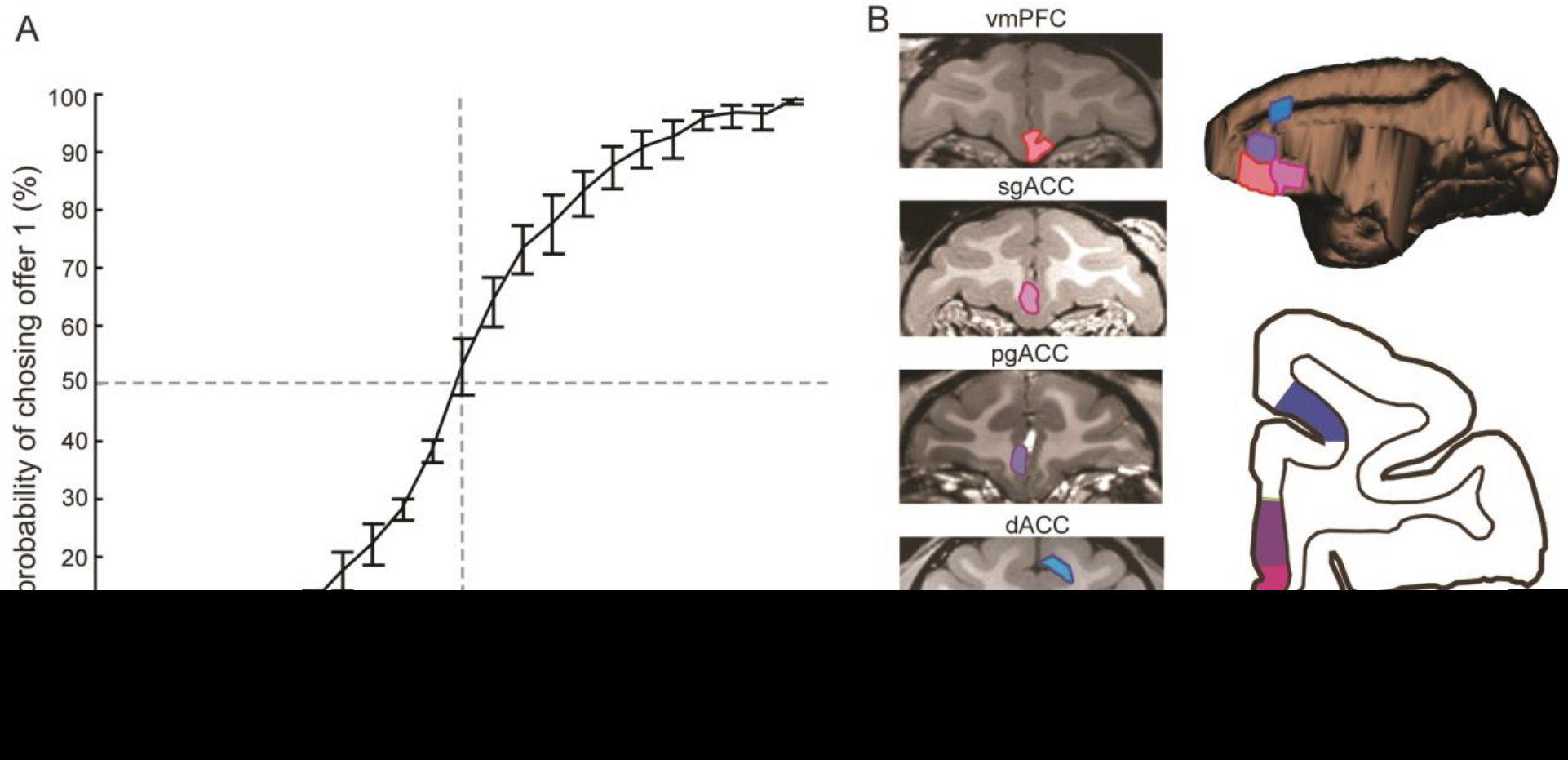
Target brain areas and behavior. A. Average behavior across all subjects and recording sessions. Error bars reflect the standard error across sessions. B. MRI scans of the brain areas targeted for recordings. A single representative subject is chosen for each area even though two subjects were recorded for each. The right panels denote the targeted brain areas on both transverse and coronal planes.

### Firing rates in all regions encode values of offers

We recorded neuronal activity from four brain regions: ventromedial prefrontal area 14 (vmPFC), subgenual anterior cingulate area 25 (sgACC), pregenual anterior cingulate area 32 (pgACC), and dorsal anterior cingulate area 24 (dACC). The data from pgACC have not been previously published. Some data from vmPFC, sgACC, and dACC have been previously published, although the key analyses here are all new (Azab and Hayden, 2017, 2018, and 2020; Strait et al., 2014, 2015, and 2016). We collected these recordings from 4 subjects (B, H, V & J see **Methods and Figure 2B**). For each area, we recorded from two subjects, although we did not use the same subjects for all areas. We did not observe marked behavioral differences across subjects and therefore did not expect nor observe qualitative differences between subjects. We collected data from dACC and sgACC in the *token risky choice task*; we collected data from vmPFC and pgACC in the *risky choice task* (**Figure 1A-B**). The basic format for each of the tasks during the selected time period for analysis, from within each trial, are essentially the same. We do not believe the small differences between the two tasks influenced the results we present here.

### Offer encoding latency does not differ between areas

First, we confirmed our hypothesis that there would be no differences between areas in stimulus (i.e. the offer) response latencies. For both offer 1 and offer 2, we computed the latency of neural responses (see **Methods**). We defined latency as the time elapsed from the onset of the offer stimulus until firing rates in the respective epoch reached their maximum within the epoch. We used a 4 (area) x 2 (offer 1 or 2) ANOVA to test for differences. Neither the main effect of area (F = 1.96, *p* = 0.297) nor offer number (F = 5.04, *p* = 0.110) was statistically significant (**Figure 3A**). Neuronal responses to the onset of offer 1 reached maximum firing (spikes per millisecond) in epoch 1 after an average of 258.55 ms from the offer onset (vmPFC = 265.6 ms; sgACC = 242.5 ms; pgACC = 252 ms; dACC = 274.1 ms). Neuronal responses in epoch 2 reached their maximum, on average, 245.04 ms after the onset of offer 2 (vmPFC = 241.8 ms; sgACC = 242.2 ms; pgACC = 245.6 ms; dACC = 250.5 ms).

**Figure 3.**
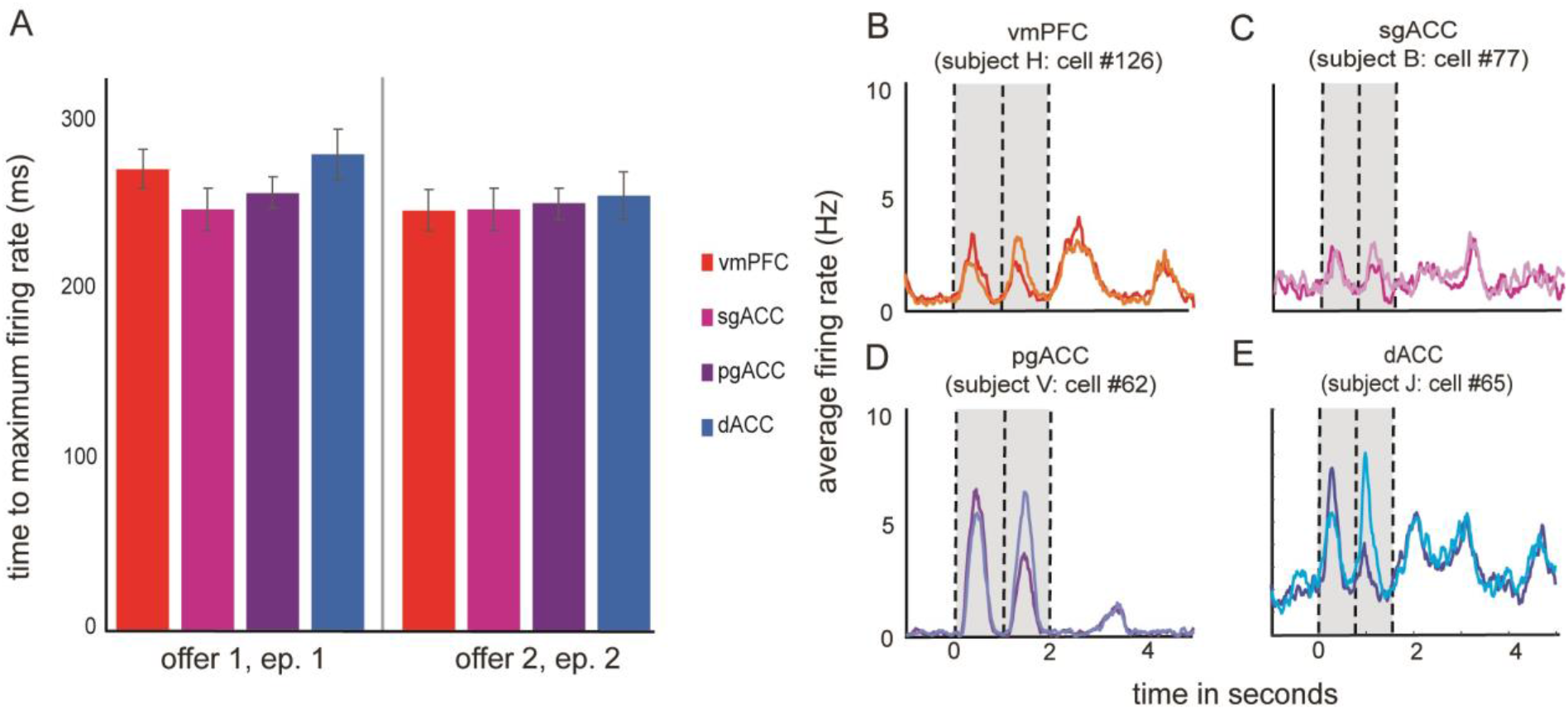
Response latency and traces of average firing rates across all trials drawn from responses of single sample neurons. A. Average latency to maximal firing rates in response to the onset of both offer 1 and offer 2. Error bars indicate the standard error across trials. B-E. Peri-stimulus time histogram responses of sample neurons with firing rates that are significantly correlated with the expected values of both offers. Traces are grouped by trials on which the value of either offer 1 (darker color) or offer 2 (lighter color) was larger. Traces are smoothed, for display, with a 200 ms sliding boxcar. Average firing rates are computed in spikes per second. The onset of offer 1 is set to time 0. The vertical lines indicate the start times of periods in the trial (the onset of offer 1, offer 2, and fixation).

We examined the proportion of neurons in each region selective for the value of offer 1 during epoch 1. **Figure 3B-E** shows the average firing rate responses of an example neuron from each of the four regions. In all four regions, a significant proportion of neurons encoded the value of offer 1 during the offer 1 epoch (vmPFC: 16.03%; sgACC: 10.27%; pgACC: 10.98%; dACC: 26.36%; *p* < 0.01 in all areas, binomial test). Likewise, neurons in three of the four regions encoded the value of offer 2 during epoch 2 (vmPFC: 14.1%; sgACC: 7.53%; pgACC: 10.98%; dACC: 14.73%; *p* < 0.01 in all areas except sgACC (trending at *p* = 0.063), binomial test). Finally, neurons encoded the value of offer 1 during epoch 2 (i.e. working memory for value, vmPFC: 9.62%; sgACC: 10.27%; pgACC: 9.41%; dACC: 14.73%; *p* < 0.01 in all areas, binomial test; **Figure 4**).

**Figure 4.**
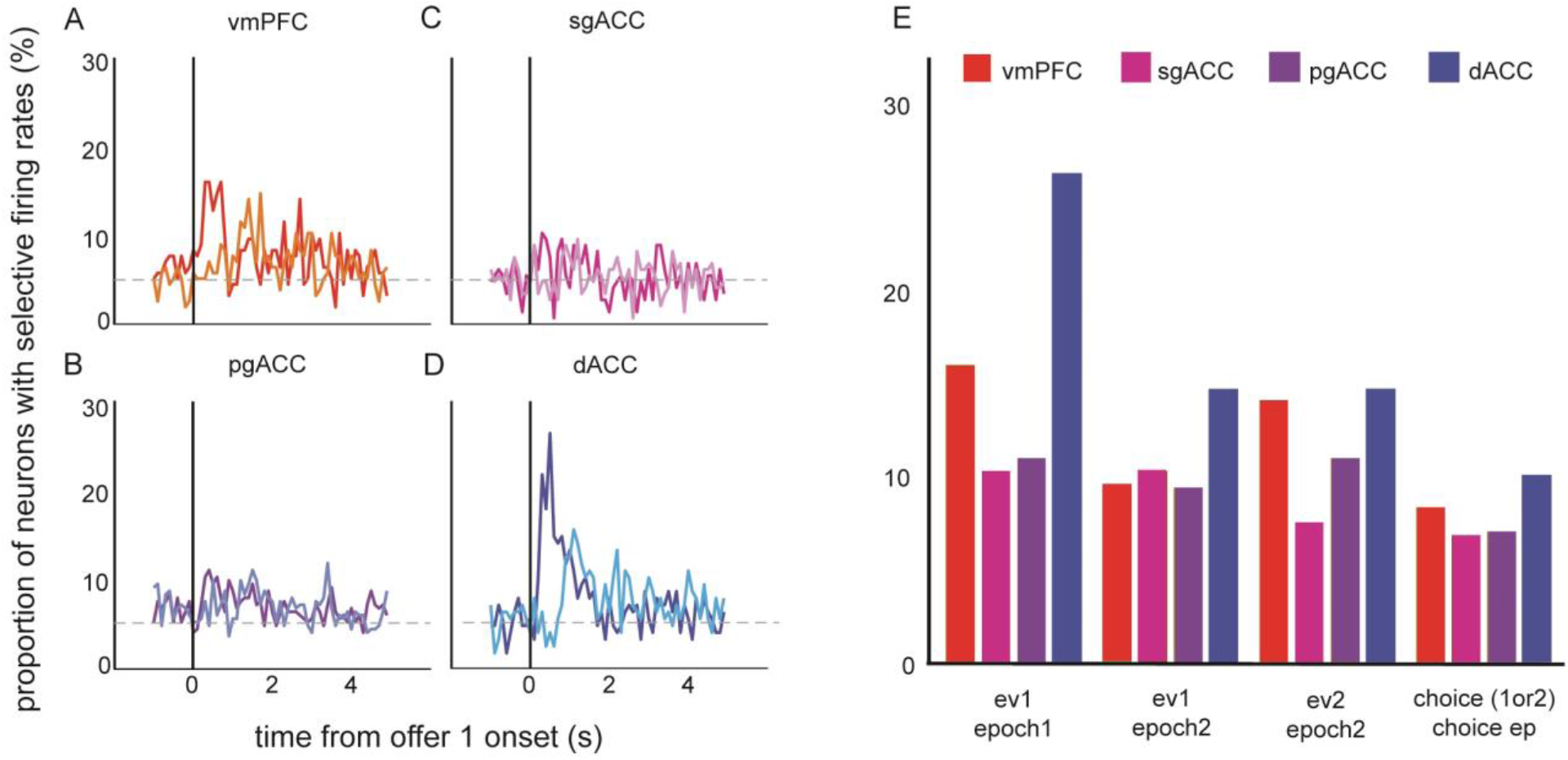
Selectivity of firing rates to offers and choice. A-D. The proportion of neurons that have firing rates that are correlated with the expected value of offer 1 (darker color) and offer 2 (lighter color) across a six second period during the trial. The onset of offer 1 is set to time 0. The vertical line indicates the onset of offer 1. E. A summary of the proportion of neurons selective to offers and choice, within given epochs. Each bar indicates the proportion within a given brain area.

### Putative signatures of choice process are found in all four regions

#### Feature Integration

We next asked whether each brain region contains a value signal that reflects the integration of the two features that determine value: probability and magnitude (Azab and Hayden, 2020). For each brain region, we computed regression weights for each neuron’s normalized (z-scored) firing rates for the two variables. We then examined how those variables related to each other across the population. A positive correlation between regression coefficients indicates that both offer features are encoded using a correlated coding scheme. In other words, it indicates that the population of neurons has thrown out information about the details of the components and has begun to compute an integrated value signal (Blanchard et al., 2015; Azab and Hayden, 2020). We observed a significant positive relationship in all regions (*p* < 0.001 in all regions; **Figure 5A**). This result indicates that feature integration is not a unique feature of any region, but instead, is broadly shared across the medial PFC.

**Figure 5.**
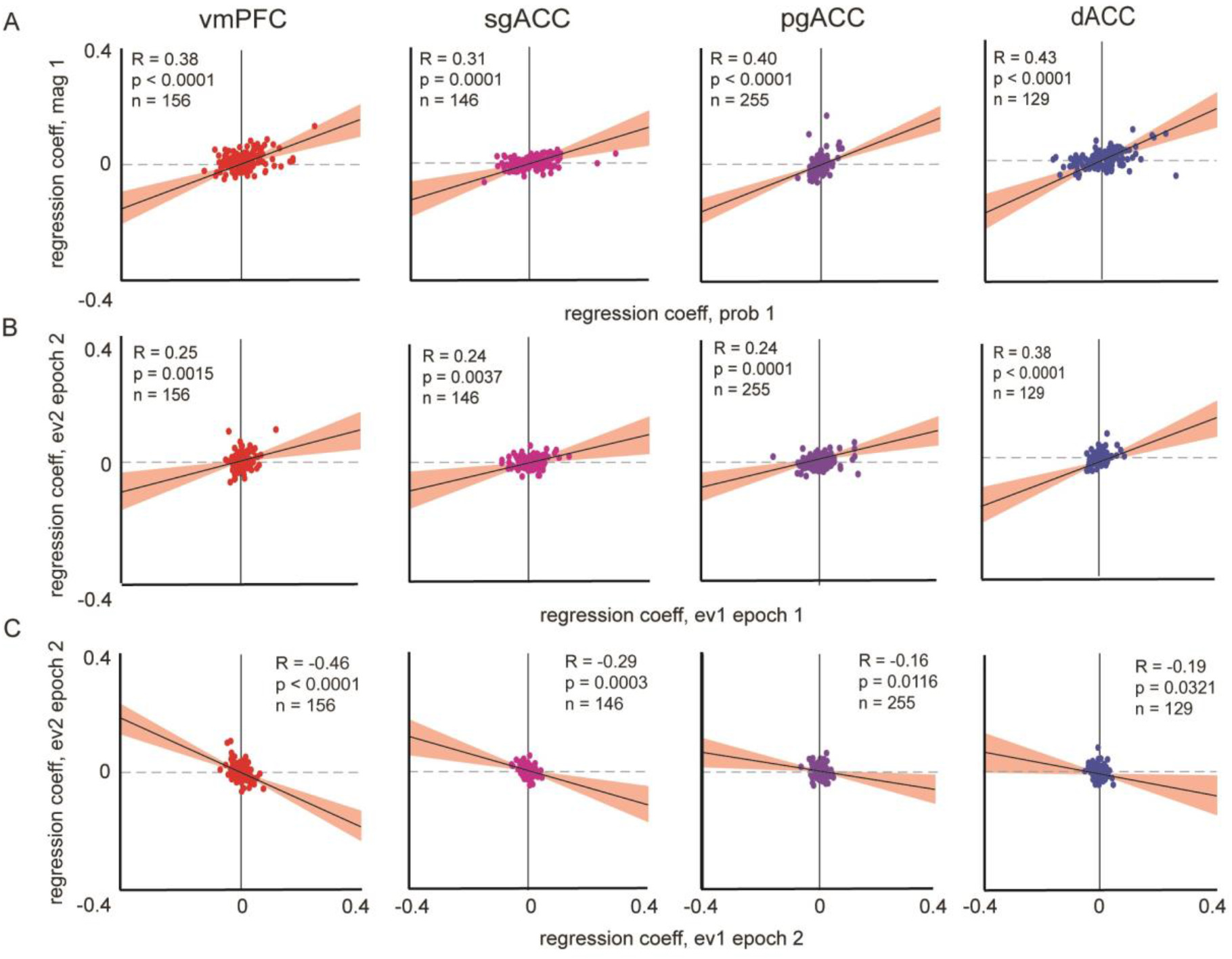
Economic choice functions. A. Feature integration. From left to right: Scatter plots of coefficients from regressing normalized epoch 1 firing rates on the probability of offer 1, against the regression coefficients from the magnitude of offer 1. The diagonal black line indicates the slope of the correlation between regression coefficients. The red ribbons indicate the 95% confidence intervals. B. Attentional alignment. Scatter plots are of coefficients from regressing normalized epoch 1 firing rates on expected value of offer 1, against the regression coefficients from epoch 2 firing rates on expected value of offer 2. C. Mutual inhibition. Scatter plots are of coefficients from regressing normalized epoch 2 firing rates on the expected value of offer 1, against the regression coefficients from epoch 2 firing rates on the expected value of offer 2.

#### Attentional alignment

When attention shifts from one option to another, it is possible that the same population of neurons encodes the new option in the same manner as it encoded the first one, like neurons in visual cortex do for visual stimuli (Hayden and Moreno-Bote, 2018; see also McGinty et al., 2016; Xie et al., 2018). In other words, neurons may act as a *flexible filter* for value; we have called this principle attentional alignment (as opposed to a labeled line code for value, Hayden and Moreno-Bote, 2018). We next asked whether attentional alignment is a principle shared in all four of our regions. To do so, we regressed normalized firing rates from epoch 1 onto the expected value of offer 1 and regressed normalized firing rates from epoch 2 on the expected value of offer 2. We then correlated these resulting coefficients. A positive correlation is evidence for attentional alignment. All four regions exhibited significant (*p* < 0.01) positive correlations (**Figure 5B**).

#### Mutual inhibition

We next asked if there was evidence for direct comparison between offers by principle of mutual inhibition (Strait et al., 2014; Azab and Hayden, 2017). We regressed normalized firing rates from epoch 2 on offer 1 onto responses from epoch 2 on offer 2. If the encoding for value 1 and value 2 during the same epoch are anti-correlated, then the encoding of 1 value comes at the expense of the other. it is an indication of the direct comparison of offers and thus a signal of choice. All recorded regions exhibited significant (*p* < 0.05) negative correlations (**Figure 5C**) between regression weights. This result indicates that all four regions show evidence of value comparison via mutual inhibition.

#### Overlapping neuronal populations

Next, we asked if there were distinct or shared populations of neurons encoding the previously described functions. We repeated the integration, alignment, and inhibition analyses (i.e. the correlations of select regression weights), but now with absolute (i.e. unsigned) values of the regression weights. We have previously shown that such correlations indicate shared or overlapping functional populations (Blanchard et al., 2018). If the (unsigned) strength of encoding of one offer (or feature) is positively correlated with the degree of encoding of the other offer, then the populations associated with encoding the two variables overlap more than expected by chance. We found a significant positive correlation (vmPFC: R = 0.31; sgACC: R = 0.31; pgACC: R = 0.44; dACC: R = 0.41; *p* < 0.001, all areas) between unsigned betas for the integration function (epoch 1 firing rates regressed on offer 1 magnitude and on offer 1 probability). We also found a significantly positive correlation between unsigned betas for all areas (vmPFC: R = 0.34; sgACC: R = 0.23; pgACC: R = 0.33; *p* < 0.01), except for dACC (R = 0.17, *p* = 0.058), representing alignment (epoch 1 firing rates regressed on expected value 1 and epoch 2 regressed on expected value 2). Finally, vmPFC (R = 0.29) and pgACC (R = 0.25) showed significantly positive correlations (*p* < 0.001 in both cases) between unsigned betas for inhibition (epoch 2 firing rates on the expected value of each offer). These results indicate that, in most areas and across functions, encoding is mostly supported by the same, or at least overlapping, sets of neurons.

### Intrinsic timescales are longest at the top of the anatomical hierarchy

These results demonstrate broadly overlapping functions across regions. We next asked whether there is evidence of hierarchy. We first considered intrinsic timescales. Intrinsic timescales are a population-level statistic describing fluctuations in a neuronal signal that are agnostic to the task and corresponding variables (Murray et al., 2014). Murray and colleagues (2014) proposed that intrinsic fluctuations are a function of increased modulatory strength. They suggested that increased modulation is due to increased recurrent network activity, which in turn increases along a hierarchy. Thus, longer intrinsic timescales would be indicative of increased modulatory strength and, therefore, a higher position along the hierarchy.

We estimated and compared the intrinsic timescales of each recorded region from a temporal decay function. We used the decay function to fit the autocorrelation of pre-trial spike data across a range of lags (**see Methods**). We found an increase of intrinsic timescale that seemed to map a medial prefrontal hierarchy onto a rough ventrodorsal gradient (vmPFC: 109.8 ms; sgACC: 152.76 ms; pgACC: 321.85 ms; dACC: 446.51 ms; **Figure 6**). We confirmed a positive monotonic relationship by correlating the intrinsic timescales with the observed order (1-4) across the areas. The results showed a significant positive correlation between increasing order across the four areas, and the increasing intrinsic timescale (R = 0.98, *p* = 0.022, Pearson’s correlation; R = 1, *p* = 0.042, Spearman’s correlation).

**Figure 6.**
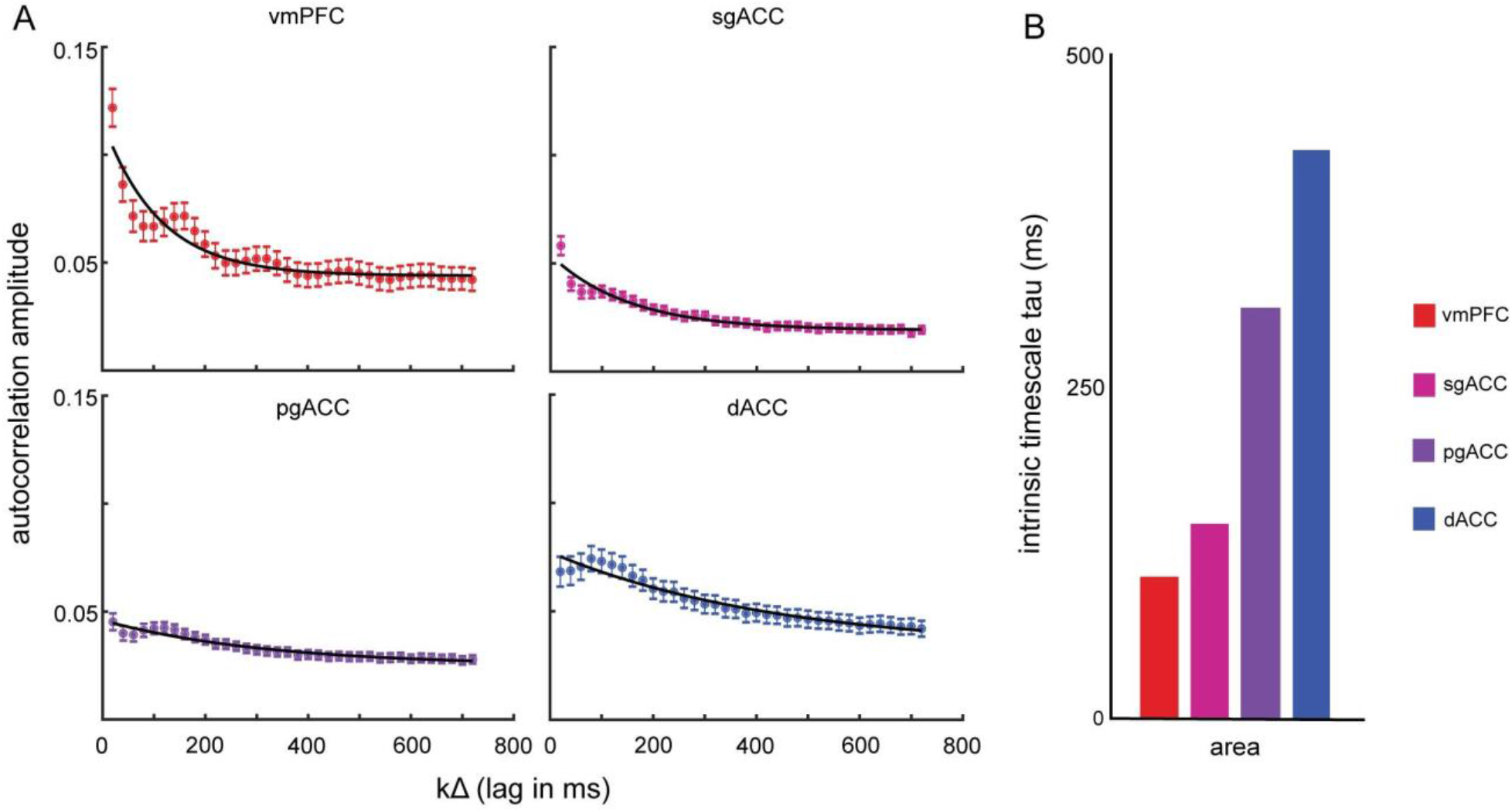
Intrinsic timescales. A. Spike-count autocorrelations calculated at increasing lags for each of the four brain areas. The decay of autocorrelation amplitude with lag was fitted by an exponential decay function with offset (black line). Error bars indicate the variance in autocorrelation amplitude across neurons at a given lag. B. A summary table of the intrinsic timescales, extracted from the exponential decay function. The figure demonstrates a smooth increase of intrinsic timescale along a hierarchy.

### Decoding accuracy supports a clear functional hierarchy

We hypothesized that the accuracy with which expected value and choice can be decoded from firing rate patterns should increase along the observed anatomical hierarchy. Thus, we trained and cross-validated a linear classifier to decode seven binary labels (specifically: high/low expected values for both offers and the difference between them, offer position, choice, and chosen side) from firing rates.

We first looked at how accurately offer 1 value could be decoded from firing rates in epoch 1 (**Figure 7A**). The classifier decoded whether offer 1 on each trial had greater or lesser value than the mean offer value across trials significantly better than chance (*p* < 0.0001, binomial test) for all regions: vmPFC (61.4%), sgACC (63.1%), pgACC (69.8%), and dACC (73.7%). Notably, decoding accuracy increases with hierarchical order (R = 0.98, *p* = 0.022, Pearson’s correlation; R = 1, *p* = 0. 042, Spearman’s correlation).

**Figure 7.**
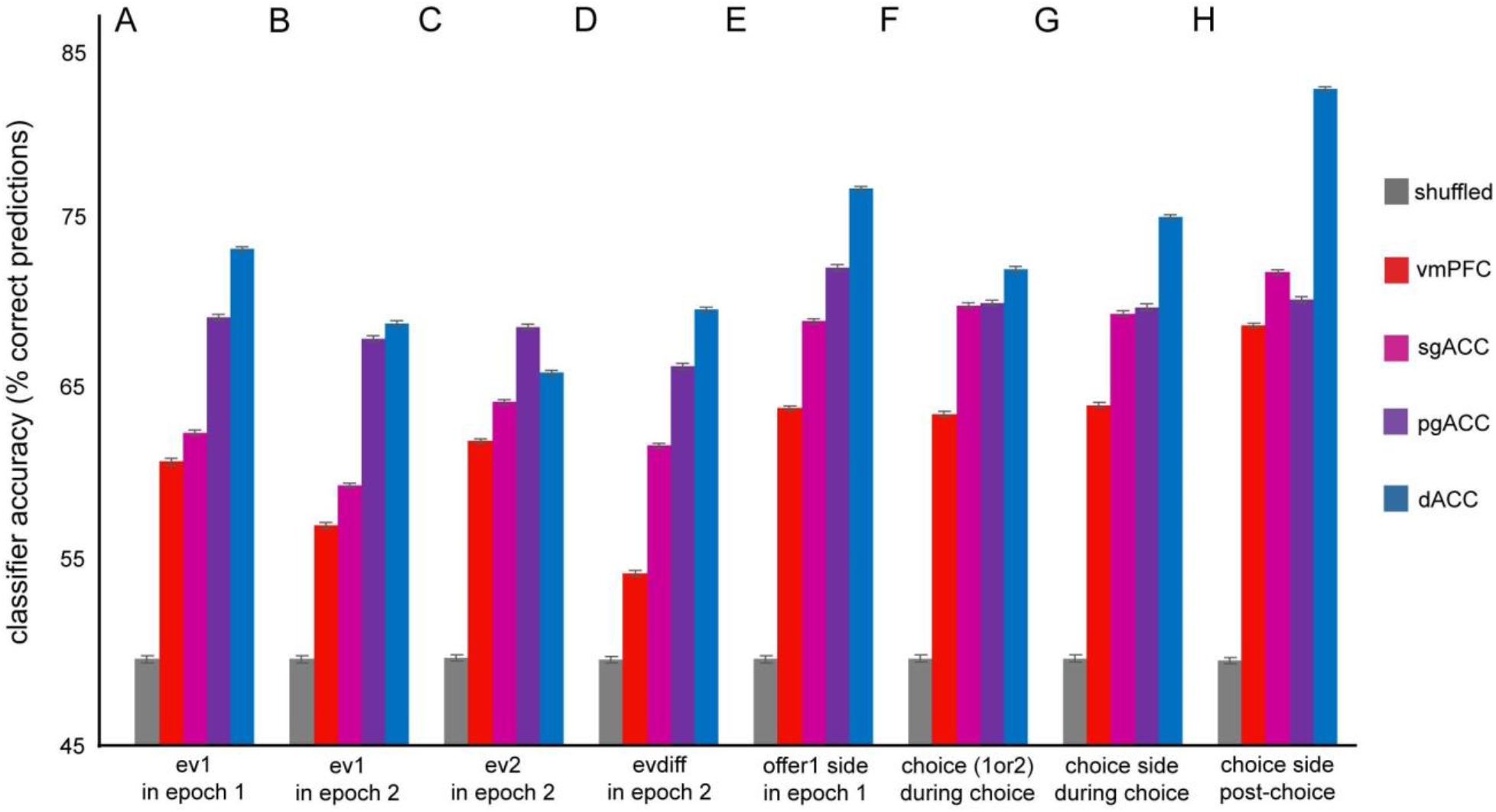
Decoding analysis. A-H. Summary of classification accuracies for each of the given labels. In a given epoch, a linear classifier was trained to identify the value of a binary label (indicated on the X-axis). The accuracy of the trained model was tested by cross-validation, to predict the label value on novel data (indicated on the Y-axis). Accuracy of a model trained on randomly shuffled data are indicated by the grey bar. Error bars represent the standard error over the variance across cross-validations.

We then looked at how accurately offer 1 value could be decoded from firing rates in epoch 2 (a putative signature of working memory for value; Kennerley and Wallis, 2009; **Figure 7B**). The classifier decoded whether offer 1 on each trial had greater or lesser value than the mean offer value across trials significantly better than chance (*p* < 0.001, binomial test) for all regions: vmPFC (57.7%), sgACC (60.1%), pgACC (68.5%), and dACC (69.4%). As above, decoding accuracy increases with hierarchical order (R = 0.95, *p* = 0.049, Pearson’s correlation; R = 1, *p* = 0. 042, Spearman’s correlation).

We next looked at how accurately offer 2 value could be decoded from firing rates in epoch 2 (**Figure 7C**). The classifier decoded whether offer 2 on each trial had greater or lesser value than the mean offer value across trials significantly better than chance (*p* < 0.0001, binomial test) for all regions: vmPFC (62.6%), sgACC (64.9%), pgACC (69.2%), and dACC (66.6%). We determined that decoding accuracy did not significantly increase with the previously observed hierarchical order (R = 0.75, *p* = 0.25, Pearson’s correlation; R = 0.8, *p* = 0.167, Spearman’s correlation). While this pattern was not significant, there still appears to be a general increase in decoding accuracy with hierarchical position.

We next looked at how accurately the difference between offer values could be decoded from firing rates in epoch 2 (**Figure 7D**). (This analysis would not make sense in epoch 1 because the subject did not know both values yet). The classifier decoded whether offer difference on each trial had greater or lesser value than the mean offer value across trials significantly better than chance (*p* < 0.001, binomial test) for all regions: vmPFC (54.9%), sgACC (62.3%), pgACC (67%), and dACC (70.2%). Decoding accuracy increases with hierarchical order (R = 0.98, *p* = 0.017, Pearson’s correlation; R = 1, *p* = 0. 042, Spearman’s correlation).

### Hierarchical organization of coding of spatial information

We looked at how accurately the position of offer on the monitor could be decoded from firing rates in epoch 1 (**Figure 7E**). The classifier decoded whether offer 1 on each trial was on the right or left (*p* < 0.0001, binomial test) for all regions: vmPFC (64.5%), sgACC (69.5%), pgACC (72.6%), and dACC (77.2%). Decoding accuracy increases with hierarchical order (R = 0.996, *p* = 0.004, Pearson’s correlation; R = 1, *p* = 0. 042, Spearman’s correlation).

Next, we looked at how accurately the chosen offer (offer 1 or 2) could be decoded from firing rates in choice epoch (**Figure 7F**). Note that this variable is orthogonal to offer side, since we observed essentially no spatial biases in choice. The classifier decoded choice on each trial (*p* < 0.0001, binomial test) for all regions: vmPFC (64.2%), sgACC (70.5%), pgACC (70.6%), and dACC (72.5%). We determined that decoding accuracy trended toward an increase with hierarchical order (R = 0.896, *p* = 0.104, Pearson’s correlation; R = 1, *p* = 0. 042, Spearman’s correlation).

We also looked at how accurately the position of the chosen offer on the monitor could be decoded from firing rates in the choice epoch (**Figure 7G**). The classifier decoded whether the chosen side on each trial was on the right or left (*p* < 0.0001, binomial test) for all regions: vmPFC (64.7%), sgACC (70%), pgACC (70.4%), and dACC (75.6%). We determined that decoding accuracy increases with hierarchical order (R = 0.96, *p* = 0.041, Pearson’s correlation; R = 1, *p* = 0. 042, Spearman’s correlation).

We also looked at how accurately the position of the chosen offer on the monitor could be decoded from firing rates in the post-choice epoch (Tsujimoto et al., 2009; **Figure 7H**). The classifier decoded whether the chosen side on each trial was on the right or left (*p* < 0.0001, binomial test) for all regions: vmPFC (69.3%), sgACC (72.4%), pgACC (70.8%), and dACC (83%). We determined that decoding accuracy does not significantly increase with hierarchical order (R = 0.82, *p* = 0.179, Pearson’s correlation; R = 0.8, *p* = 0.167, Spearman’s correlation). While this pattern was not significant, there still appears to be a general increase in decoding accuracy with hierarchical position.

## DISCUSSION

Here we examined neuronal correlates of multiple elements of economic choice in four medial prefrontal cortex regions. Confirming and extending our previous results, we find that these regions show largely similar value-related signals (Strait et al., 2014; Azab and Hayden, 2018). Indeed, by none of the measures we chose did these regions differ qualitatively. This result suggests that the regions do not have conspicuous qualitative differences along the dimensions we studied, but leaves open the possibility that they differ quantitatively. Our major novel finding is that, by several measures, the regions appear to be organized hierarchically. First, eight basic task variables are consistently more decodable later in the hierarchy. These include both abstract (economic) and spatial variables. Second, intrinsic timescale is longer later in the hierarchy. Overall, our results are consistent with the idea that the four regions serve as part of a roughly ventral-to-dorsal functional gradient that gradually transforms neural encodings (Yoo and Hayden, 2018).

The idea that prefrontal regions have a largely hierarchical organization was pioneered by Fuster, who proposed a functional gradient from the sensory to the motor areas, and involving “association cortex” between them (Fuster, 2001). Although he (like many subsequent thinkers) was mainly focused on the lateral prefrontal cortex, the same logic may extend to medial areas. However, the most logical organization of such areas is not obvious, either anatomically or functionally. There are many possibilities. Primarily using anatomical connectivity patterns, Price and colleagues classify all four of our recorded regions in his “medial network,” which they propose are responsible for visceromotor functions, and contrasting with the “orbital network,” responsible for sensory functions (Carmichael and Price, 1994; Carmichael and Price, 1996; Ongur and Price, 2000).

Based on cytoarchitecture and laminar connectivity patterns, Barbas and Pandya (1989) take a somewhat different view. For them, areas 25, 24, and 32, as relatively undifferentiated cingulate cortex, are all placed in a similar, low position in a mediodorsal hierarchy. Area 14, split between the mediodorsal and basoventral trends, occupies a somewhat higher position in the hierarchy. Our results (although we interpret them differently with respect to a hierarchy) are not necessarily inconsistent with such a framework, as increased decodability and timescales may simply be a hallmark of less-differentiated PFC regions.

Alternatively, topology would suggest possible *ventrodorsal* (14→25→32→24) or *genu-adhering* (25→14→32→24) hierarchies. The first one is consistent with the idea that OFC (Price’s orbital network) serves as the entryway for economic information to the prefrontal cortex, and area 14 as its next station (Rushworth et al., 2011; Noonan et al., 2010). Our work supports the ventrodorsal hypothesis most strongly, thereby offering the first electrophysiological evidence for one specific medial prefrontal hierarchy. One prediction of this hierarchy is that medial area 9 (dorsomedial prefrontal cortex) should be one step above the recorded areas in our analyses (cf. Schall et al., 2002). We might also expect that sensory choice information is received by orbitofrontal cortical area 13, and then relayed to the medial prefrontal cortex; thus we would expect area 13 to be below the recorded areas in this hierarchy, to have shorter intrinsic timescales, and to have less decodable information.

Our results suggest that these four regions have largely overlapping functions in the domain of economic choice. Notably, our results do not imply that these regions have identical functions, nor that their differences are solely quantitative. Indeed, there is plentiful evidence that these regions have important qualitative differences (Bush et al, 2000; Vogt and Laureys, 2005; Van Holstein and Floresco, 2020). To give an example, in a social aggression paradigm, activation of the ventral medial prefrontal cortex correlates with skin conductance response, perhaps reflecting its strong interactions with hypothalamus and periaqueductal gray, while activation of the dorsal medial prefrontal cortex is more cognitive in nature (Lotze et al., 2007). Our results do not challenge or invalidate such categorical functional differences. Rather, they suggest that these regions have qualitative differences in some domains and quantitative differences in at least one domain, the domain of economic choice. Indeed, our results do point to a potential limitation to much of traditional functional neuroanatomy. Much of that work is focused exclusively on identifying the unique contributions of particular regions. While that work is critically important, it necessarily ignores the kinds of brain functions that are not uniquely implemented by specific regions. We believe that economic choice is one such function (Yoo and Hayden, 2018).

Nonetheless, a broad reading of the electrophysiological literature highlights that many functions have traces that are quite similar in multiple regions (Hunt and Hayden, 2017). Many scholars draw a distinction between sensory areas, for which a strong modularity case can be made, and “association areas”. For example, as Prinz (2006) points out, even Fodor, a great advocate of modularity, was more willing to consider distributed function outside of sensory and motor regions (Fodor, 1983). Likewise, Uttal (2001) identifies Olds’ work on classical (trace) conditioning (1972), which shows that correlates of trace conditioning can be found in nearly every part of the rat brain.

## METHODS

### Surgical procedures

The University Committee on Animal Resources at the University of Rochester and University of Minnesota approved all animal procedures. Animal procedures were designed and conducted in compliance with the Public Health Service’s *Guide for the Care and Use of Animals*. Four male rhesus macaques (*Macaca mulatta*) served as subjects for both tasks. A small prosthesis head fixation was used. Animals were habituated to laboratory conditions and then trained to perform oculomotor tasks for liquid rewards. We place a Cilux recording chamber (Crist Instruments) over the area of interest (see *Behavioral tasks* for breakdown). We verified positioning by magnetic resonance imaging with the aid of a Brainsight system (Rogue Research). Animals received appropriate analgesics and antibiotics after all procedures. Throughout both behavioral and physiological recording sessions, we kept the chamber with regular antibiotic washes and we sealed them with sterile caps.

### Recording sites

We approached our brain regions through standard recording grids (Crist Instruments) guided by a micromanipulator (NAN Instruments).

We defined **vmPFC 14** as lying within the coronal planes situated between 29 and 44 mm rostral to the interaural plane, the horizontal planes situated between 0 and 9 mm from the brain’s ventral surface, and the sagittal planes between 0 and 8 mm from the medial wall (**Figure 2B**). These coordinates correspond to area 14 in Paxinos et al. (2008).

We defined **sgACC 25** as lying within the coronal planes situated between 24 and 36 mm rostral to interaural plane, the horizontal planes situated between 17.33 and 25.12 mm from the brain’s dorsal surface, and the sagittal planes between 0 and 5.38 mm from medial wall (**Figure 2B**). Our recordings were made from central regions within these zones, which correspond to area 25 in Paxinos et al. (2008).

We defined **pgACC 32** as lying with the coronal planes situated between 30.90 and 40.10 mm rostral to the interaural plane, the horizontal planes situated between 7.30 and 15.50 mm from the brain’s dorsal surface, and the sagittal planes between 0 and 4.5 mm from the medial wall (**Figure 2B**). Our recordings were made from central regions within these zones, which correspond to area 32 in Paxinos et al. (2008).

We defined **dACC 24** as lying within the coronal planes situated between 29.50 and 34.50 mm rostral to interaural plane, the horizontal planes situated between 4.12 to 7.52 mm from the brain’s dorsal surface, and the sagittal planes between 0 and 5.24 mm from medial wall (**Figure 2B**). Our recordings were made from central regions within these zones, which correspond to area 24 in Paxinos et al. (2008).

We confirmed recording location before each recording session using our Brainsight system with structural magnetic resonance images taken before the experiment. Neuroimaging was performed at the Rochester Center for Brain Imaging on a Siemens 3T MAGNETOM Trio Tim using 0.5 mm voxels. We confirmed recording locations by listening for characteristic sounds of white and gray matter during recording, which in all cases matched the loci indicated by the Brainsight system with an error of ~1 mm in the horizontal plane and ~2 mm in the z-direction.

### Electrophysiological techniques

Either single (FHC; starting impedance 4 ㏁) or multi-contact electrodes (V-Probe, Plexon) were lowered using a microdrive (NAN Instruments) until waveforms between one and three neuron(s) were isolated. Individual action potentials were isolated on a Plexon system (Plexon, Dallas, TX) or Ripple Neuro (Salt Lake City, UT). Neurons were selected for study solely on the basis of the quality of isolation; we never preselected based on task-related response properties. All collected neurons for which we managed to obtain at least 300 trials were analyzed; no neurons that surpassed our isolation criteria were excluded from analysis.

### Eye-tracking and reward delivery

Eye position was sampled at 1,000 Hz by an infrared eye-monitoring camera system (SR Research). Stimuli were controlled by a computer running Matlab (Mathworks) with Psychtoolbox and Eyelink Toolbox. Visual stimuli were colored rectangles on a computer monitor placed 57 cm from the animal and centered on its eyes (Fig. 1*A*). A standard solenoid valve controlled the duration of juice delivery. Solenoid calibration was performed daily.

### Behavioral tasks

Four monkeys performed in two different tasks with the same basic structure. For the neuronal recordings in vmPFC, *subjects B* and *H* performed the *risky choice task*; and for dACC and sgACC, *subjects B* and *J* performed the *token risky choice task* (Fig. 2*C*). Both tasks made use of vertical rectangles indicating reward amount and probability. We have shown in a variety of contexts that method provides reliable communication of abstract concepts such as reward, probability, delay, and rule to monkeys (Hayden et al., 2009; Blanchard and Hayden 2014; Sleezer et al., 2016; Mehta et al., 2019).

### Risky choice task (vmPFC and pgACC 32)

All tasks were based on a standardized general structure for gambling tasks (Hayden et al, 2010; Heilbronner and Hayden, 2013; Heilbronner and Hayden, 2016; Heilbronner, 2017). The task presented two offers on each trial. A rectangle 300 pixels tall and 80 pixels wide represented each offer (11.35° of visual angle tall and 4.08° of visual angle wide; Fig. 2*A*). Two parameters defined gamble offers, *reward size* and *probability*. Two portions divided each gamble rectangle, one red and the other either grey, blue, or green. The size of the color portions signified the probability of winning a small (125 μl), medium (mean 165 μl), or large reward (mean 240 μl), respectively. We drew a uniform distribution between 0 and 100% for these probabilities. Red colored the rest of the bar; the size of the red portion indicated the probability of no reward. Offer types were selected at random with a 43.75% probability of blue (medium magnitude) gamble, a 43.75% probability of green (high magnitude) gambles, and a 12.5% probability of gray options (safe offers).

On each trial, one offer appeared on the left side of the screen and the other appeared on the right. We randomized the sides of the first and second offer (left and right). Each offer appeared for 400 ms and was followed by a 600-ms blank period. After the offers were presented separately, a central fixation spot appeared and the monkey fixated on it for 100 ms. Following this, both offers appeared simultaneously and the animal indicated its choice by shifting gaze to its preferred offer and maintaining fixation on it for 200 ms. Failure to maintain gaze for 200 ms did not lead to the end of the trial but instead returned the monkey to a choice state; thus monkeys were free to change their mind if they did so within 200 ms (although in our observations, they seldom did so). Following a successful 200-ms fixation, the trial immediately resolved the gamble and delivered the reward. We considered trials that took ~7 s as inattentive trials and we did not include them in the analyses (this removed ~1% of trials). Outcomes that yielded rewards were accompanied by a visual cue: a white circle in the center of the chosen offer. All trials were followed by an 800-ms intertrial interval with a blank screen.

### Token risky choice task (sgACC 25 and dACC 24)

Another similarly structured gambling task, where gambles each had two potential outcomes, wins or losses in terms of “tokens” displayed on screen as cyan circles. A small reward (100 μl) was administered concurrently with gamble feedback on each trial, regardless of gamble outcome. Trials in which the monkey accumulated six or more tokens triggered an extra “jackpot” epoch in which a very large reward (300 μl) was administered (Fig. 2*C*).

### Behavioral analysis

To confirm the statistical validity of the behavioral results, we first identified the chosen offer (first or second) for each trial. We then calculated the proportion of trials, for each recording session, in which the offer with the higher value was chosen. We then analyzed the vector of choice proportions using a 1-sample t-test, to determine if the average proportion of greater value choices was statistically different from zero.

### Reuse of data

Some of these data were previously published (vmPFC dataset in Strait et al. 2014; sgACC and dACC data sets in Azab and Hayden, 2017; data from pgACC have not been previously published).

### Statistical methods

We constructed peristimulus time histograms by aligning spike rasters to the presentation of the first offer and averaging firing rates across multiple trials. We calculated firing rates in 20-ms bins but we generally analyzed them in longer (500 ms) epochs. For display, we smoothed peristimulus time histograms using a 200-ms running boxcar. Some statistical tests of neuron activity were only appropriate when applied to single neurons because of variations in response properties across the population. In such cases, a binomial test was used to determine if a significant portion of single neurons reached significance on their own, thereby allowing conclusions about the neural population as a whole.

### Offer encoding latency

We computed an average latency score for each area and both offers. First, we isolated firing rates for both epochs, as each was binned to the onset of one of the offers. Each epoch consisted of a 500 ms window constituted by 20 ms bins. Next, we calculated the average firing rate for each neuron, for each 20 ms bin, across trials. Then, for each neuron, we determined how much time (in ms) passed for a given neuron to reach its peak firing rate for the epoch. Finally, we calculated the average latency to peak firing rate across all neurons in the region.

### Intrinsic timescales

To measure intrinsic timescales, we followed similar steps described in Murray, et al. (2014). We isolated a 2-second time window preceding the onset of offer 1, to remain independent of trial variables. Using a 20 ms sliding window, we then computed the autocorrelation for the 2 second window with a given lag *k*Δ between time *i* and time *j,* where *k*Δ= |*i - j*|. The lag ranged from 20ms and 720ms. We then determined that the autocorrelation decay in each structure could be well-fit by an exponential decay function:

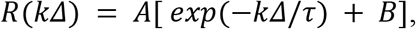

where *A* = the amplitude of the autocorrelation, *k*Δ= the lag, τ = intrinsic timescale, and *B* = the offset to account for long timeframes outside of the measured window. This formula follows what was described by Murray, et al. 2014.

### Decoding analysis

We built a pseudo-population of pseudo-trials. First, we isolated each epoch and collapsed the firing rates for each trial into an average for the 500 ms period. Then, we separated the data set for each neuron by the given label (1st offer and second; left offer and right; offer 1 position left or right; offer 2 position left or right; offer 1 value higher or lower than the mean value; and offer 2 value higher or lower than the mean value). We randomly selected 1000 samples for each neuron resulting in 2 n X 1000 matrices (one for each label level), where n represented the number of neurons recorded from each region. This constituted the pseudo-population or pseudo-trials. To execute the decoder, each matrix was split in half and concatenated with the half from the other label. We used one of these matrices to train a binary support vector machine, the other was used for cross-validation. We used the trained model to predict the binary label for each pseudo-trial in the cross-validation set. We then compared predicted outcome to the known choice outcome and an accuracy rate was calculated across pseudo-trials. This process was repeated 100 times for each target structure and for each of the given labels and epochs to get a distribution of accuracy rates. Thus, the standard error of the mean, used in displaying the error bars, represents the standard error over the variance of the cross-validations. Additionally, the exact process was repeated on randomly shuffled data, to confirm that expected prediction accuracy was 50% when randomized.

## Acknowledgements

We thank Meghan Castagno Pesce, Marc Mancarella, Caleb Strait and Tommy Blanchard for assistance with data collection, and the rest of the Hayden/Zimmermann lab for valuable discussions. This research was supported by a National Institute on Health grant R01 DA038106 (to BYH) a R01 MH118257 (to SRH), a National Institute on Drug Abuse grant P30 DA048742-01A1 (to BYH, SH and JZ), a National Institute for Biomedical Imaging Grant P41 EB027061 (to BYH and JZ), and a UMN AIRP award (to BYH, SH and JZ).

## Notes

### Competing Interest Statement

The authors have declared no competing interest.

